# Anterior Nucleus of Thalamus Gates Progression of Mesial Temporal Seizures by Modulating Thalamocortical Synchrony

**DOI:** 10.1101/2020.09.17.301812

**Authors:** Ganne Chaitanya, Adeel Ilyas, Emilia Toth, Diana Pizarro, Kristen Riley, Sandipan Pati

## Abstract

The anterior nucleus of the thalamus (ANT) mediates cortical-subcortical interactions between the limbic system and is hypothesized to facilitate the early organization of temporal lobe seizures. We set out to investigate the dynamic changes in synchronization parameters between the seizure onset zone (SOZ) and ANT during seizure stages (pre-onset to post-termination) in seven patients (n=26 seizures) with drug-resistant nonlesional temporal lobe epilepsy. Using local field potentials recorded directly from the limbic system and the ANT during stereoelectroencephalography, we confirm that the onset of mesial temporal lobe seizure is associated with increased thalamocortical network excitability and phase-amplitude coupling. The increase in thalamocortical phase synchronization preceded seizure onset, thereby suggesting that the early organization of temporal lobe seizures involves the integration of the ANT within the epileptic network. Towards seizure termination, there is a significant decrease in thalamic excitability, thalamocortical synchronization, and decoupling, thereby suggesting a breakdown in thalamocortical connectivity. A higher disease burden is significantly correlated with increased synchronization between the ANT and epileptic networks. Collectively, the results elucidate mechanistic insights and provide the temporal architecture of thalamocortical interactions that can be targeted in the rational designing of closed-loop seizure abortive interventions.

**Highlights:**

- Anterior nucleus of thalamus is coactivated at the onset of temporal lobe seizures
- Increase thalamocortical synchronization and excitability is observed at seizure onset
- Seizure termination is characterized by a breakdown in thalamocortical connectivity
- Increased seizure burden affects thalamocortical synchronization

## 1. Introduction

Pathologically synchronized neural activity within local and long-range networks characterizes state transitions as seizures evolve from interictal to onset, propagation, and termination stages(Jiruska et al., 2013)(Kramer et al., 2010)(Netoff and Schiff, 2002). Understanding the nature of these state transitions is critical in developing effective antiseizure therapies(Sohal and Sun, 2011)(Yu et al., 2018). Temporal lobe epilepsy (TLE) is a common form of drug-resistant focal epilepsy that is surgically remediable with an anterior temporal lobectomy (ATL). However, seizures recur in 40-65% of patients following resection(Josephson et al., 2013)(Bell et al., 2009). Emerging evidence suggests that recurrent seizures contribute to the reorganization of networks involving temporal and extratemporal regions, including the thalamus(He et al., 2017)(Keller et al., 2015). Neural interactions within the aberrant thalamocortical network can sustain seizures in preclinical models of focal epilepsy(Paz et al., 2013)(Feng et al., 2017)(Langlois et al., 2010). However, the pathophysiologic mechanisms by which thalamocortical interactions support seizures in focal epilepsies are poorly understood.

The thalamus serves as a crucial hub within the thalamocortical network and modulates information processing between cortical regions(Jones, 2001)(Steriade, 1997). Observations from preclinical studies suggest that cortical feedback to the thalamus modulates the thalamic neuronal firing patterns and generates synchronous oscillatory activity that conjoins information flow between cortical regions(Destexhe, 2000)(Bal et al., 2000). Limbic seizures recruit the thalamus and modulate the degree of thalamocortical synchronization, which may be variable at the onset but is typically heightened near seizure termination(Guye et al., 2006)(Feng et al., 2017)(Aracri et al., 2018). In particular, the anterior nucleus of the thalamus (ANT) mediates cortical-subcortical interactions between the limbic system and brainstem and facilitates seizure propagation(Takebayashi et al., 2007)(Sherdil et al., 2019). In several preclinical studies, chemical or electrical modulation of the ANT disrupted the progression of limbic seizures, thereby establishing its causal role in ictogenesis(Covolan et al., 2014)(Sherdil et al., 2019)(Bittencourt et al., 2010). Indeed, headway has been made in the development of thalamocortical neuromodulation therapies in focal epilepsies. For example, the approved ANT DBS provides chronic open-loop stimulation during the interictal state and, at best, yields a modest decrease in seizure frequency(Salanova et al., 2015). However, the mechanism of how the thalamic modulation can perturb seizure is poorly understood, and changes in synchronization are one proposed candidate mechanism(Yu et al., 2018).

We set out to investigate the dynamic changes in synchronization parameters between the seizure onset zone (SOZ) and ANT during peri-ictal states in a cohort with drug-resistant TLE. Local field potentials (LFPs) were recorded directly from multiple nodes within the limbic system, including the SOZ and the ANT during stereo EEG (SEEG) investigation for localization of suspected TLE. In a previous study, we observed an increase in population activity within the ANT at the onset and propagation of temporal lobe seizures(Pizarro et al., 2019)(D. et al., 2018). In the present study, we examined the changes in intrinsic excitability of the ANT and coupling between the ANT and SOZ as mesial temporal onset seizures evolve from pre-seizure to posttermination stages. Coupling of oscillations between lower and higher frequency bands has been proposed to coordinate neural information flow at multiple spatial scales, including the thalamocortical network(FitzGerald et al., 2013)(Ibrahim et al., 2018). We hypothesized that the ANT would be coactivated at the onset of temporal lobe seizures. With the seizure progression, there would be changes in intrinsic thalamic excitability that modulate intracortical information flow via changes in thalamocortical coupling. In this way, we provide not only mechanistic insights into the role of the ANT in TLE but also define the temporal architecture of thalamocortical neural communications during seizure stages.

## 2. Methods

### 2.1 Ethics statement

The study was approved by the Institutional Review Board of the University of Alabama at Birmingham (IRB170323005). Written informed consent was obtained to perform research recordings from the thalamus. Through a meticulous planning and consenting process as detailed previously, adults with drug-resistant suspected TLE were educated that the trajectory of one of the clinical depth electrodes sampling the frontal operculum and insula would be modified to additionally sample the ANT(Chaitanya et al., 2020). This approach would obviate the need for additional electrode placement for research purposes, thereby minimizing the risk of hemorrhage. Patients were explicitly informed about the risks associated with research implantation, and our group has recently reported the consent process and safety of thalamic implantation that was accompanied by an editorial(McKhann, 2020)(Chaitanya et al., 2020).

### 2.2 Patient selection

Patients with suspected TLE underwent stereotactic electroencephalography (SEEG) when noninvasive studies were discordant or inconclusive for localization of the seizure foci. The study inclusion criteria were: a) SOZ must include the mesial temporal structures (amygdala-hippocampus; b) absence of epileptogenic lesion in the 3-Tesla MRI brain (i.e., nonlesional TLE) to avoid structural deformity confounding the results; c) confirmed the localization of the depth electrode within the ANT, ipsilateral to SOZ; and, d) a minimum of 6-9 months follow up posttherapy. The seizure outcome was graded as per Engle classification at the last clinic follow-up(He et al., 2017)(Engel et al., 1993).

### 2.3 SEEG implantation: mapping the targets and trajectory

Our SEEG implantation strategy and accuracy have been published previously (JNS FOCUS reference). In summary, robot-assisted electrode implantation (diameter: 0.8mm, 10-16 contacts per electrode, contact length: 2mm, inter-contact distance: 1.5mm, PMT^®^ Corporation, Chanhassen, MN, and ROSA^®^ device, Medtech) of predetermined regions of interest for seizure localization was performed. The trajectory of the ANT implantation followed passing through precentral gyrus, pars opercularis frontalis, insula, and putamen before ending in the ANT(Chaitanya et al., 2020). The following imaging sequences were used to confirm the localization of electrodes: preimplantation MRI sagittal T1-weighted images (3T Philips Achieva, voxel size: 1×1×1mm, FOV: 170×256×256mm) and post-implantation CT axial images (Philips Brilliance 64 scanner, 1mm slices, in-plane resolution of 0.44×0.44mm, FOV: 228×228×265mm). ANT contacts were localized using Lead-DBS v2 software and the cortical contacts using iElectrodes(Horn et al., 2019). Preoperative MRI and postoperative CT scans were linearly co-registered using Advanced Normalization Tools followed by refinement with brain shift correction to improve the registration of subcortical structures(Jenkinson et al., 2002). Both the images were normalized to ICBM 2009b NLIN asymmetric space using the symmetric diffeomorphic image registration. The cortical regions of all the implanted SEEG contacts were confirmed with AAL2 atlas, and the thalamic nuclei were identified using the mean histological thalamic Morel’s atlas (Fig1 A-B)(Krauth et al., 2010).

**Figure 1:**
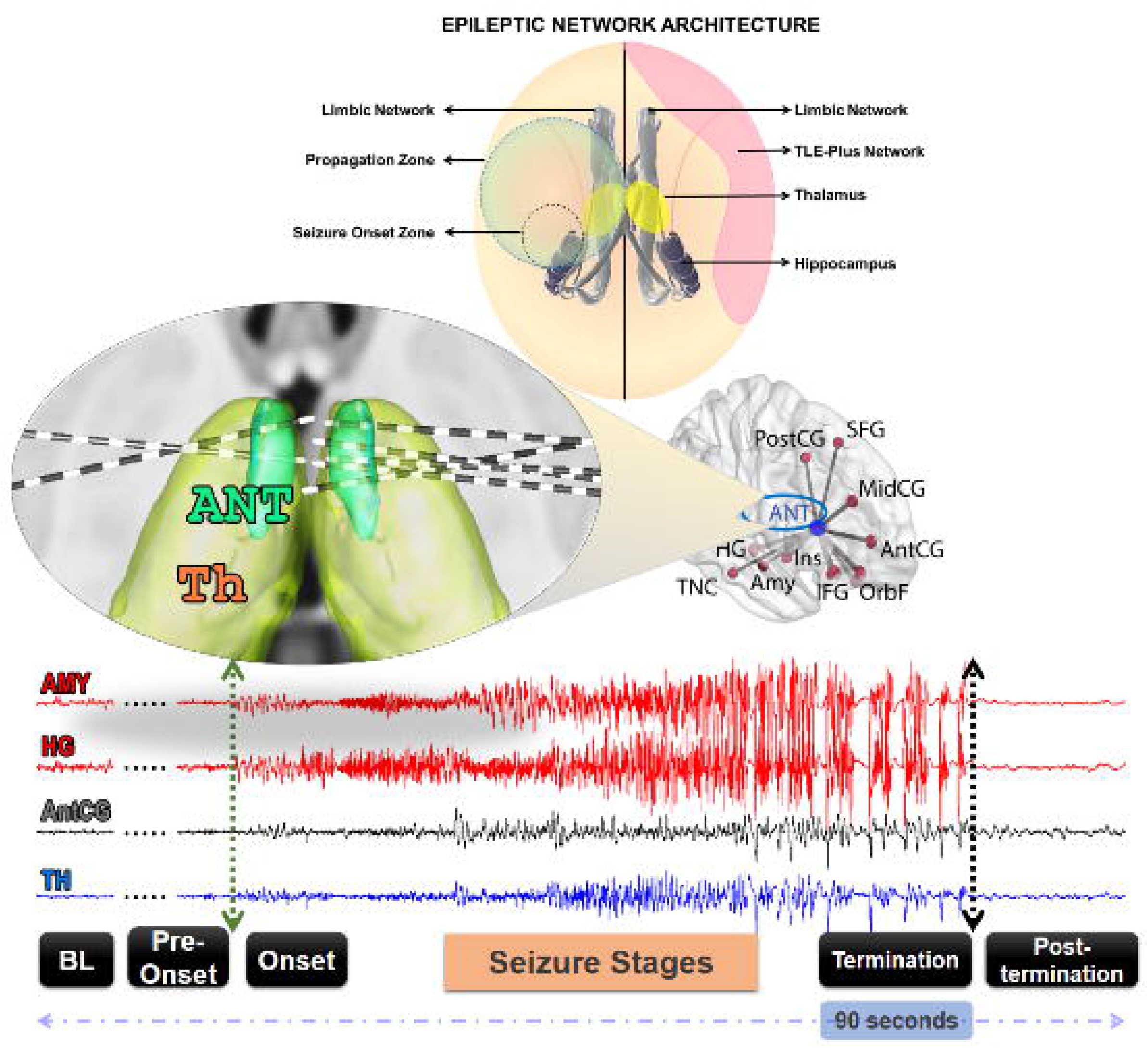
Schematic representation of regions of interest sampled with stereo-electroencephalography (AC) and an example of seizure recording (D). (A) Anatomically, all patients had certain common regions that were sampled, i.e., regions of the medial, lateral temporal lobe, and limbic network (includes orbitofrontal, insula, and ant. cingulum). Functionally, the regions were grouped per patient as - (i) seizure onset zone, (ii) propagation zone, and (iii) noninvolved regions. Collectively these three regions were labeled as ENA (epileptic network architecture). (B) Image reconstruction (reconstructed using Morel’s thalamic atlas in LeadDBS.v2) confirming the implantation in the anterior nucleus of thalamus (ANT) in 7 patients. (C) The commons regions sampled across all patients. (D) An example of seizure starting in the hippocampal (HG)-amygdala (AMY) complex and subsequently spreading to the thalamus (TH). The peri-ictal period was divided into four stages: Baseline (BL), Pre-Onset, Onset, Termination, and Post-termination. OrbF-orbitofrontal; AntCG-anterior cingulate gyrus; MidCG-midcingulate gyrus; PostCG-posterior cingulate gyrus; SFG-superior frontal gyrus; IFG-inferior frontal gyrus; Ins-insula; TNC-temporal neocortex; ANT-anterior nucleus of thalamus.

### 2.4 SEEG data analysis

SEEG was sampled at 2048Hz using Natus Quantum (Natus Medical Incorporated, Pleasanton, CA). Seizures were annotated by a board-certified epileptologist (SP) and were categorized into subtypes: focal aware seizures (FAS), focal impaired awareness seizures (FIAS), and focal to bilateral tonic-clonic seizures (FBTCS)(Fisher et al., 2017). To eliminate potentially confounding large-scale artifacts and common source noise on the reference channel, we calculated an along-the-chain bipolar montage. The signals were subsequently notch-filtered at 60Hz. Remnants of structured artifacts running across multiple channels were excluded using independent component analysis (Jader algorithm, EEGLAB)(Jung et al., 2000)(Makeig et al., 1996) after the data were downsampled to 512Hz and the sorted independent components were manually scrutinized for structured artifacts and subtracted. The artifact-free components were back-projected and used to reconstruct the SEEG data free of artifacts.

### 2.5 Identifying and defining regions of interest

Although the implantation strategy was driven by clinical necessity and was individualized for each patient based on the underlying clinical hypothesis, all patients had certain commonly sampled anatomic regions. These regions included the hippocampus, amygdala, temporal pole, superior and middle temporal gyri, orbitofrontal gyrus, cingulate gyrus, insula, and inferior frontal gyrus (Fig1 A,C)(Barba et al., 2016). Functionally, these temporal and extratemporal lobe regions were further categorized into the following:

1. Seizure Onset Zone (SOZ): This was defined as the set of cortical structures with high epileptogenicity and early onset ictal activity. These channels were initially identified by clinical consensus after reviewing of semiology, imaging findings, and SEEG results. Subsequently, these initially selected channels were narrowed based on the epileptogenicity index (EI). EI statistically summarizes the spectral and temporal parameters of SEEG signals and quantifies the propensity of a brain area to generate low voltage fast discharges, thereby providing a quantitative metric of epileptogenicity (Bartolomei et al., 2008). The EI was computed across all channels during seizure onset, and those with an EI > 0.2 were considered to be with the SOZ(Pizarro et al., 2019)(Roehri et al., 2018).
2. Propagation Zone (PZ): This was defined as the set of cortical structures in which the EI > 0.2 but were not within the SOZ.
3. Noninvolved zone (NIZ): This was defined as the set of cortical structures that were shared sampled across all patients, except those within the SOZ and PZ. Thus, the EI for these channels were <0.2.
4. Epileptic Network Architecture (ENA): This was defined as the union of the SOZ, PZ, and NIZ (i.e., ENA = SOZ ∪ PZ ∪ NIZ).

### 2.6 Selection of seizures for the study

Only focal seizures (FAS, FIAS) that recruited the ANT were included in this study. FBTC seizures were recorded from one subject only that precludes performing group statistics and hence excluded from the study. Ictal recruitment of the ANT was quantitatively determined using the line length (LL) algorithm(Esteller et al., 2001). The Line Length feature is a simplification of the Katz’s running fractal of a signal and is proportional to changes in amplitude and frequency variations of a seizure. The specific parameters used in this study were a moving average window of 0.25 seconds (s), with a 50% overlap. The threshold of detection was set at two standard deviations above the mean LL of a 4□minute baseline segment, and the algorithm flagged a segment if the LL was persistently above the threshold for at least 10 seconds. These quantitative results were visually inspected, and the localizations were confirmed by ascertaining the presence of unequivocal electrographic changes at seizure onset within the identified SOZ and ANT channels.

### 2.7 Identifying and defining periods of interest during a seizure

Seizure onset was defined as the earliest occurrence of rhythmic or repetitive epileptiform spikes (Fig 1D). The electrographic onset of the seizure was annotated by visual inspection by a board-certified epileptologist (SP). For each seizure, five periods of interest were selected. Baseline was a 1-minute segment that was free of artifacts and interictal spikes. The baseline was selected approximately 10 minutes prior to seizure onset. The remaining periods of interest were 4 seconds long each and consisted of (1.) Pre-onset: prior to the annotated electrographic onset of the seizure in the SOZ, (2.) Onset: immediately after the identifiable electrographic onset of the seizure, (3.) Termination: prior to electroclinical seizure offset, and (4.) Post-termination: immediately after seizure offset.

### 2.8 Estimation of power spectral density (PSD) calculation

Power spectral changes during the same time interval were computed by applying the fast Fourier transform on 4s consecutive segments with 50% overlap (i.e., Welch’s method with Hamming window) per seizure and baseline stages from both SOZ and ANT channels(Welch, 1967). The PSD values were averaged per frequency band (delta [1–4Hz], theta [4–8Hz], alpha [8–13Hz], beta [13–30Hz], gamma [30–70Hz], high gamma [70-150Hz]) and then z-normalized relative to the mean of the baseline segment to ensure comparability across the stage.

### 2.9 Estimation of the spectral exponent from the PSD

We evaluated the non-oscillatory, 1/f-like activity constituting the background of electrophysiological signals, which decays from slower to faster frequencies as per the inverse power law(Gao et al., 2017)(Voytek et al., 2015). This measure indexes a subtle balance of excitation and inhibition determined by changes in high-frequency power (e.g., gamma) in relation to the low-frequency power (e.g., delta) and is considered a surrogate of cortical excitability. It was quantified using the slope of a linear regression of the power spectral density (PSD 1-150Hz, Welch’s method, Hanning window:3s, overlap: 67%) fit to the logarithm of x and y axes (i.e., broadband spectral exponent-β)(Colombo et al., 2019). Studies have suggested that the steepening of the PSD, as measured by a more negative spectral exponent, is associated with increased synchronization of neural assemblies, increased inhibition, and dampening of propagation(Miskovic et al., 2019)(Freeman and Zhai, 2009)(Gao et al., 2017)(Miller et al., 2009). The changes in intrinsic excitability were measured in the SOZ and ANT during baseline, onset, and termination windows.

### 2.10 Estimation of thalamocortical phase synchrony

Phase lag index (PLI) estimates phase synchronization that is invariant against the presence of common source artifacts whose phase differences are likely to center around 0 and π radians(Stam et al., 2007). The absence of uniform distribution of phases between two signals implies that the two signals are in synchrony. PLI was estimated in the following way. The downsampled, ICA cleaned bipolar data of each of the four periods were divided into 2s moving windows with a 50% overlap. An index of the asymmetry of the phase difference distribution can be obtained from a time series of phase differences (Δ□ (tk), k=1 … N) estimated using Hilbert transform was estimated as:

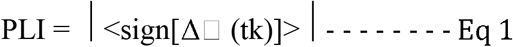

where the PLI ranges between 0 and 1. The PLI values were averaged into 2 interaction domains: (1.) widespread phase synchrony between ANT and ENA (termed global PLI [gPLI]) and (2.) localized phase synchrony between the ANT and SOZ. The PLI values were averaged across the previously mentioned frequency bands.

### 2.11 Phase amplitude coupling calculation

Whereas PLI provides information regarding phase coupling between two regions within a given frequency, phase-amplitude coupling (PAC) estimates interaction of the phase of low-frequency (LF) oscillations that modulate the amplitude of high-frequency (HF) oscillations of the two signals xLF(t) and xHF(t)(Canolty and Knight, 2010). Their complex analytic representations zLF(t) and zHF(t) are derived by means of Hilbert transform of narrowband filtered data. The envelope A(t) and instantaneous phase □(t) signal are dealt with independently. Next, the envelope of the higher-frequency oscillations AHF(t) is bandpass-filtered within the range of LF oscillations. The resulting signal is Hilbert transformed to isolate its phase-dynamics component □’(t). This reflects the modulation of HF-oscillations amplitude by the phase of the LF-oscillations. Here, the two signals were from the SOZ and the ANT. We tested the PAC in both directions, i.e., LF of SOZ modulating the HF of ANT and vice versa. Comparisons of PAC were made based on the baseline. We evaluated how the PAC changes from a pre-onset to onset and, subsequently, how the onset transitions into a termination state.

### 2.12 Statistical methods

The raw PSD values were log-transformed, and inter-stage comparisons were made with paired t-tests with a false discovery rate (FDR) correction. The interstage comparisons in the spectral exponent were also tested with paired t-tests with FDR correction and not ANOVA since we mainly wanted to report specific, individual comparisons rather than comparisons of the model as a whole. The raw PLI values were baseline normalized. Dynamical changes in PLI were tested using repeated-measures ANOVA followed by post-hoc analysis with Bonferroni correction. All the tests were modeled for multiple comparisons based on the apriori test hypothesis involving a combination of the stage of seizure, the frequency band of analysis to avoid redundancy of multiple comparisons. Stepwise multiple regression analysis was used to test the influence of long-term disease burden measures such as duration of epilepsy, seizure duration, and frequencies on thalamocortical synchrony at onset and termination. Finally, significant PAC changes consisted of values outside the mean ± 2 standard deviations of change in modulation index between any two seizure stages (e.g., onset versus termination).

## 3. Results

### 3.1 Patient cohort

The demographic details of the seven patients are described in **Table 1**. In five patients, the ANT was targetted on the right side, and, in two patients, it was targetted on the left side. None of the patients had an intracranial hemorrhage or cerebral infarction, as evaluated by post-explantation CT imaging(Chaitanya et al., 2020). All patients had seizure onset in the hippocampal-amygdala complex ipsilateral to the implanted ANT. Overall, we analyzed 26 focal seizures that met the inclusion criteria (see section 2.6).

**Table 1:**
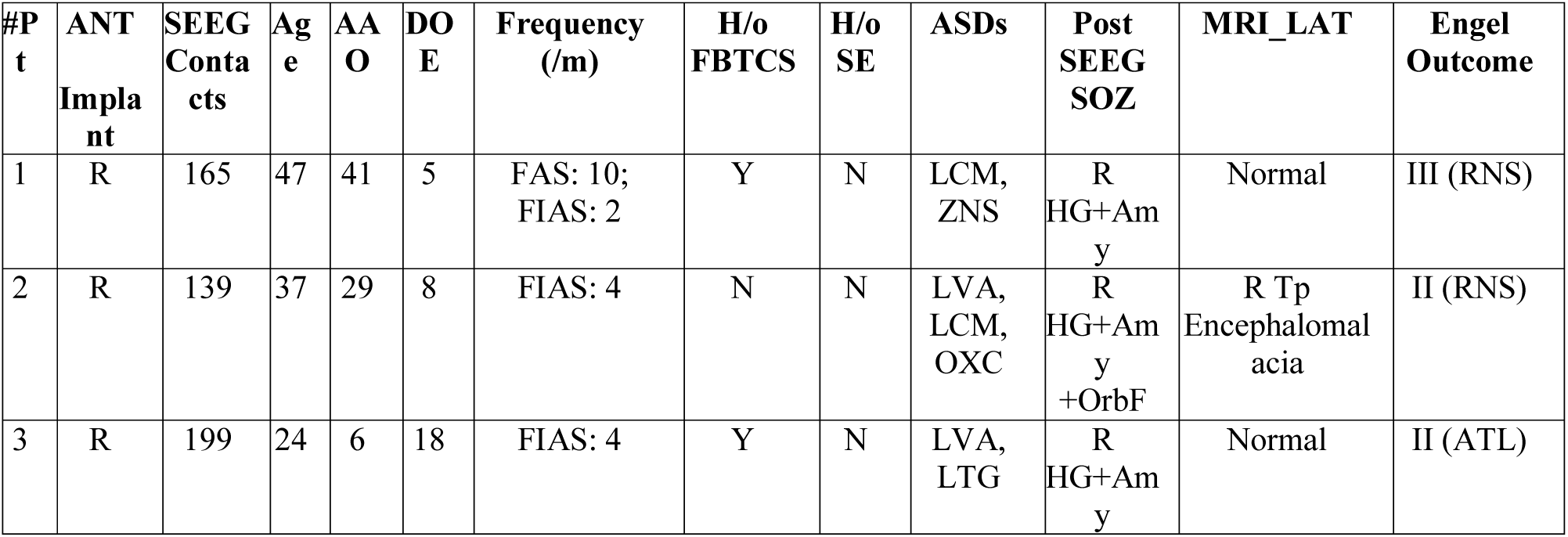
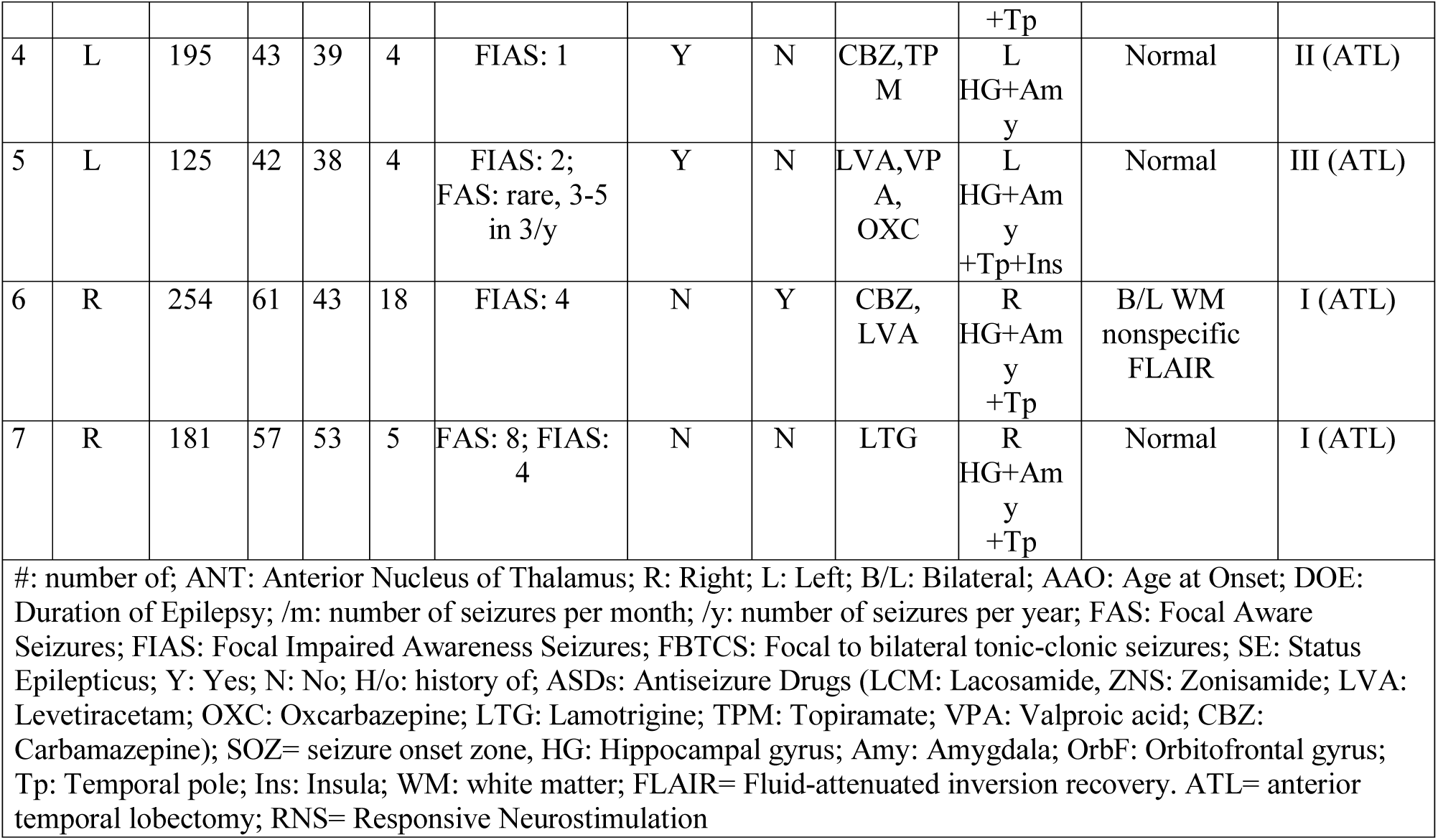
Clinico-demographic details of the patients:

### 3.2 Cortico-thalamic population activity at seizure onset and termination: changes in power spectra and spectral exponents (β) of 1/f

As anticipated, seizure onset was characterized by a significant increase in the broadband(1-150Hz) power of local population activity within the SOZ (_FDR_Ps<0.0072, ts>-3.5). Similarly, at the onset, there was an increase in the LFP broadband power in the ANT (_FDR_Ps<0.0004, ts>-3.98), although the rise in high gamma was not significant from the baseline(Fig2). At termination, the presence of continuous neural activity in both the ANT and SOZ was evident by higher power spectra, and this was increased significantly in comparison to the onset (SOZ: p=0.003, t=-2.23; ANT: p=0.0006, t=-3.81).

**Figure 2:**
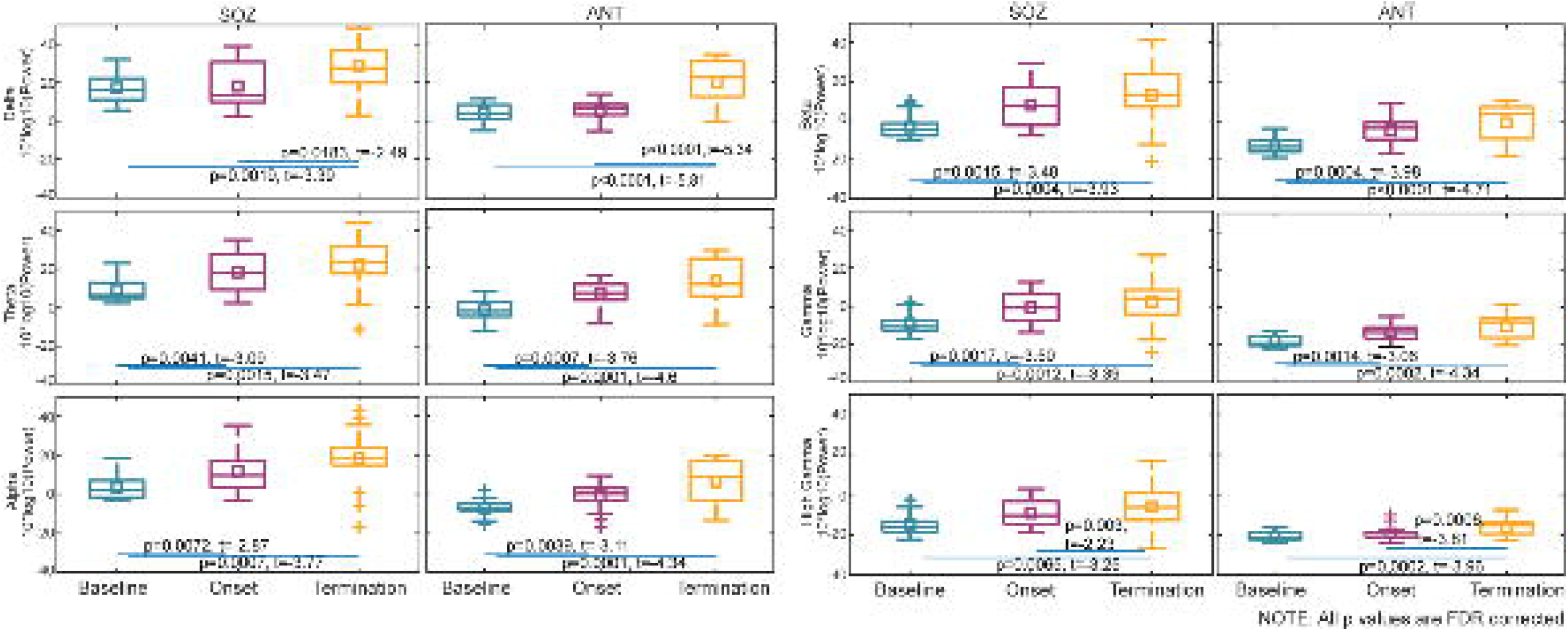
Stage-wise changes in cortico-thalamic power spectra at seizure onset and termination: Fast Fourier transform-based power-spectral analysis was performed at baseline, at seizure onset, and at the termination of both the seizure onset zone (SOZ) and the anterior nucleus of thalamus (ANT). It was evaluated across various frequency bands (delta-1-4Hz, theta(4–8Hz), alpha (8–13Hz), beta (13–30Hz), gamma (30–70Hz), high gamma (70-150Hz). The raw power values were log-transformed (log10), and inter-stage comparisons were corrected with false discovery rate correction (Pfdr was considered signifiacnt). At seizure onset ANT showed an increase broadband power from theta to low gamma range (FDRPs<0.0004, ts>-3.98). At Termination, however, the ANT showed increased high gamma power compared to onset (SOZ: p=0.003, t=-2.23; ANT: p=0.0006, t=-3.81).

Two distinct patterns of changes in the negative steepness of the power-spectral 1/f slope were observed in the SOZ and ANT (Fig3). Within the SOZ, a linear translation of the spectral exponent slope was observed. This was secondary to a broadband increase in spectral power (i.e., a relative increase in power from delta through high gamma). Within the ANT, however, a splaying pattern of the spectral exponent slopes was observed, implying an asymmetric change in spectral power between different bandwidths. Specifically, during termination, in the ANT, there was a relative increase in power in the lower frequencies (delta and theta) and increased steepness of the slope when compared with the onset(_FDR_P=0.0003, t=4.42). Such an increase in negative slope suggests decreased excitability (higher inhibition than excitation) of the ANT(Gao et al., 2017)(Colombo et al., 2019). In contrast, the spectral exponent slope of ANT at onset (compared to baseline) showed a splaying in the higher frequencies, indicating an increase in neural activities in high-frequency.

**Figure 3:**
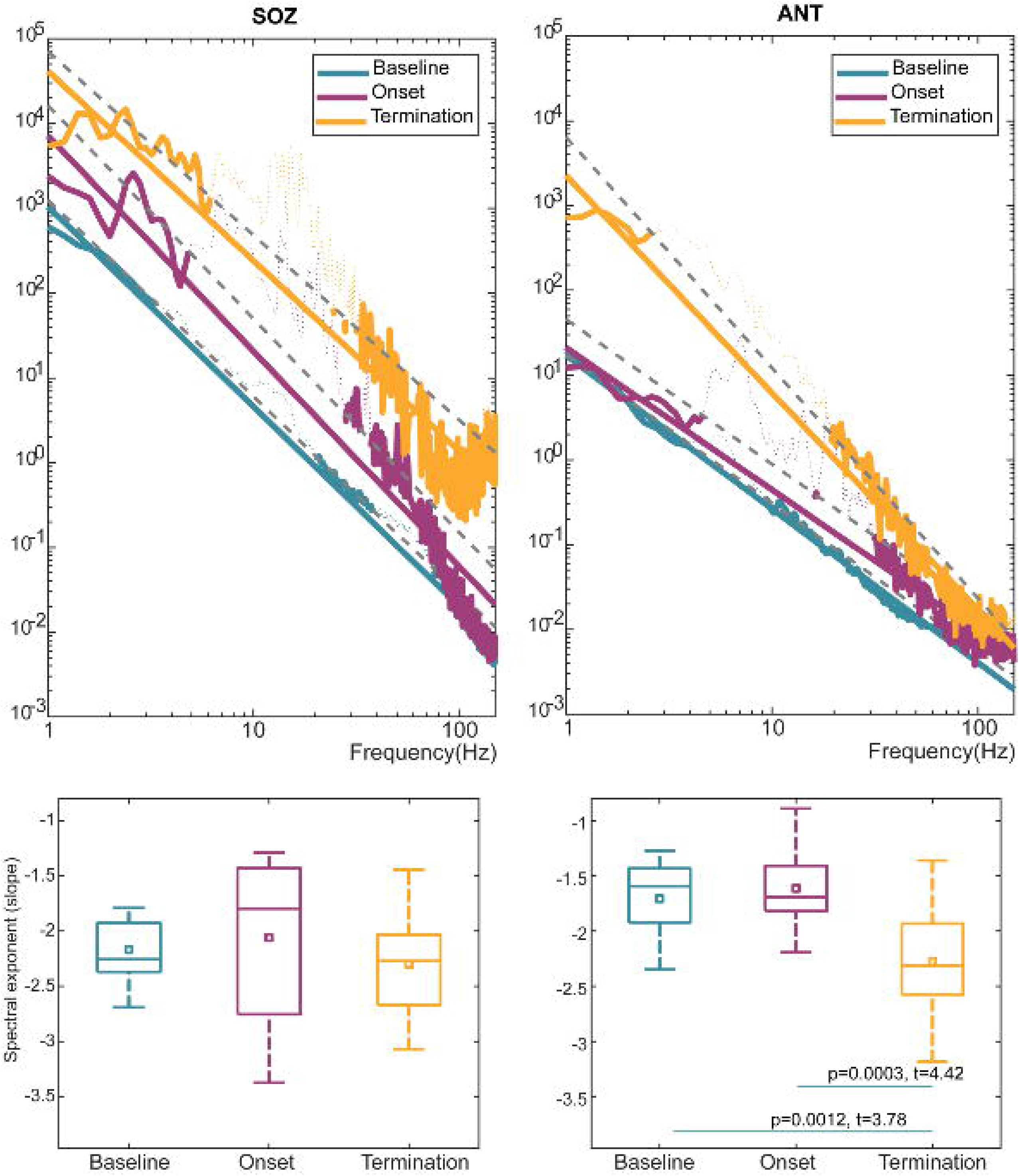
Stage-wise changes in cortico-thalamic spectral exponent as a marker of tissue excitability: Spectral exponents (β) is a surrogate marker of tissue excitability. Seizure onset zone (SOZ) and anterior nucleus of thalamus (ANT) had distinct patterns of spectral exponent change as seizure evolved from onset into termination. A broadband increase in spectral power in the SOZ resulted in a linear translation of the spectral exponent slope (i.e., a relative increase in power from delta through high gamma) between termination (yellow line) and onset (purple line). However, in ANT a splaying pattern of the spectral exponent slopes was observed, implying an asymmetric change in spectral power between different bandwidths with higher delta-theta power at termination (yellow line) compared to onset (purple line).

### 3.3 Global thalamocortical phase synchronization at seizure onset and termination

During the pre-onset stage, there was increased phase synchronization between the ANT and ENA (estimated using global PLI) in all frequencies except high gamma(Fig4). Onset was characterized by increased theta, beta, and low gamma synchrony (f’ s>11.65, df=3, _Bonferroni_P’ s<0.004). Termination was characterized by an increased delta synchrony (f=4.45, df=3, _Bonferroni_P=0.005) and a decreased theta and beta synchrony (f’ s>11.68, df=3, _Bonferroni_P’ s<3.7×10^−7^). Post-termination was characterized by a decrease in theta, alpha, and beta synchrony to baseline level (f’ s>6.76, df=3, _Bonferroni_P’ s<0.0002; all post-hocs are reported at _Bonferroni_Ps<0.05).

**Figure 4:**
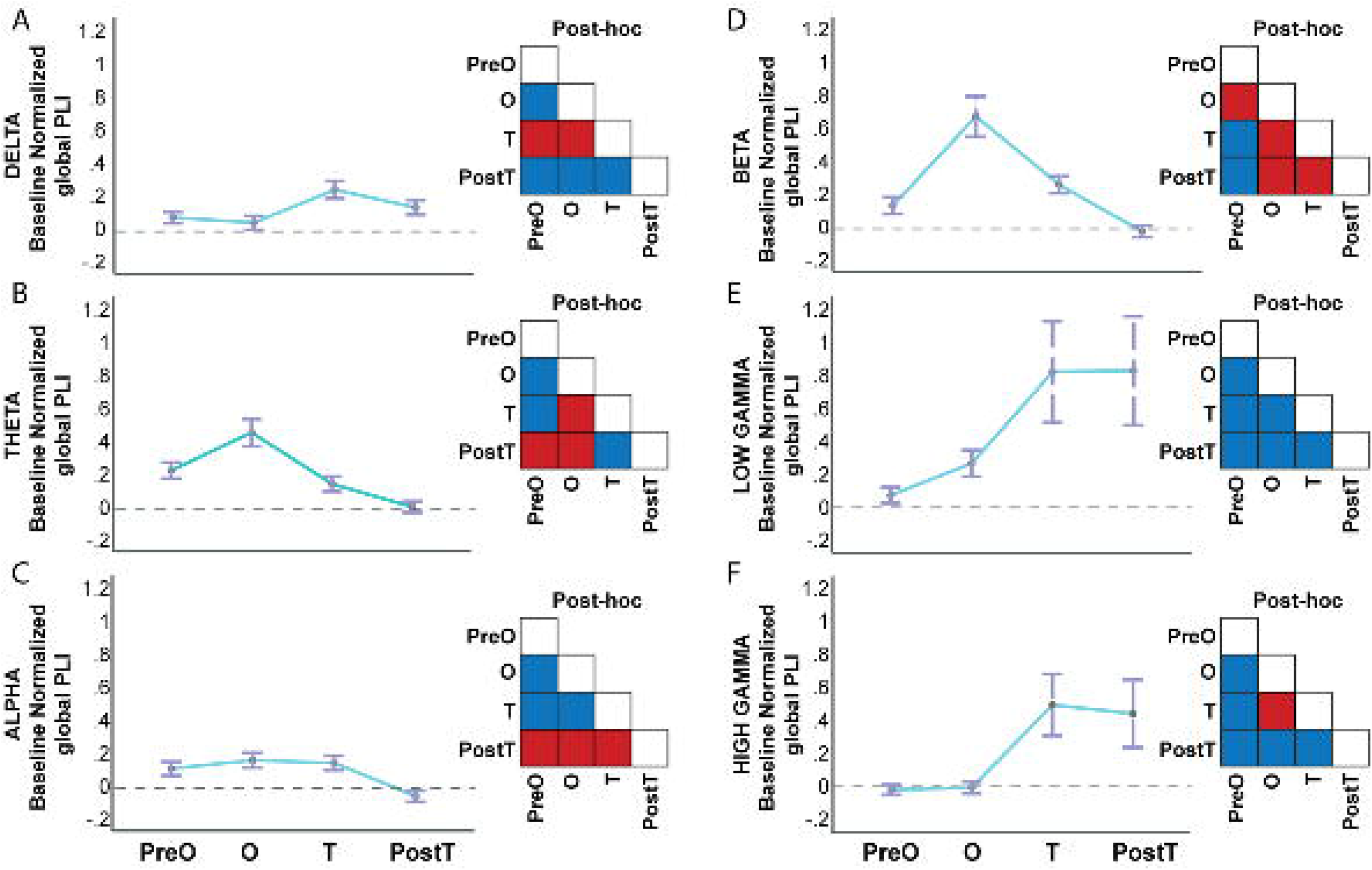
Global thalamocortical phase synchronization during seizure evolution: Global thalamocortical phase synchronization was estimated as the phase lag index (global PLI) between the anterior nucleus of the thalamus (ANT) and all the structures within the epileptic network architecture (ENA). The PLI values were baseline normalized. It was measured across the stages preonset (PreO), onset (O), Termination (T), and post-termination (PostT). (A-E) The line graphs with the error bars (mean±standard error) represent the change in phase synchrony across the different stages (preO to PostT) and frequencies (delta to high gamma). The corresponding lower-triangle insets represent inter-stage post hoc comparisons (Red – significant difference between stages and the blue no-significant difference between stages following Bonferroni correction).

### 3.4 ANT-SOZ localized phase synchronization at seizure onset and termination

We specifically evaluated dynamic change in phase synchrony between ANT and SOZ(Fig 5). The pre-onset stage was characterized by increased synchrony over a broadband of frequencies, including high gamma. At the onset, the theta and beta synchrony was highest compared to post-termination (which was comparable to baseline level of synchrony between ANT and SOZ) (f’ s>4.49, df=3, _Bonferroni_P’ s<0.006). At termination, beta synchrony was lower compared to onset but only significantly higher compared to post-termination (f=8.62, df=3, p=5.5×10^−5^)(All post-hocs are reported at _Bonferroni_Ps<0.05).

**Figure 5:**
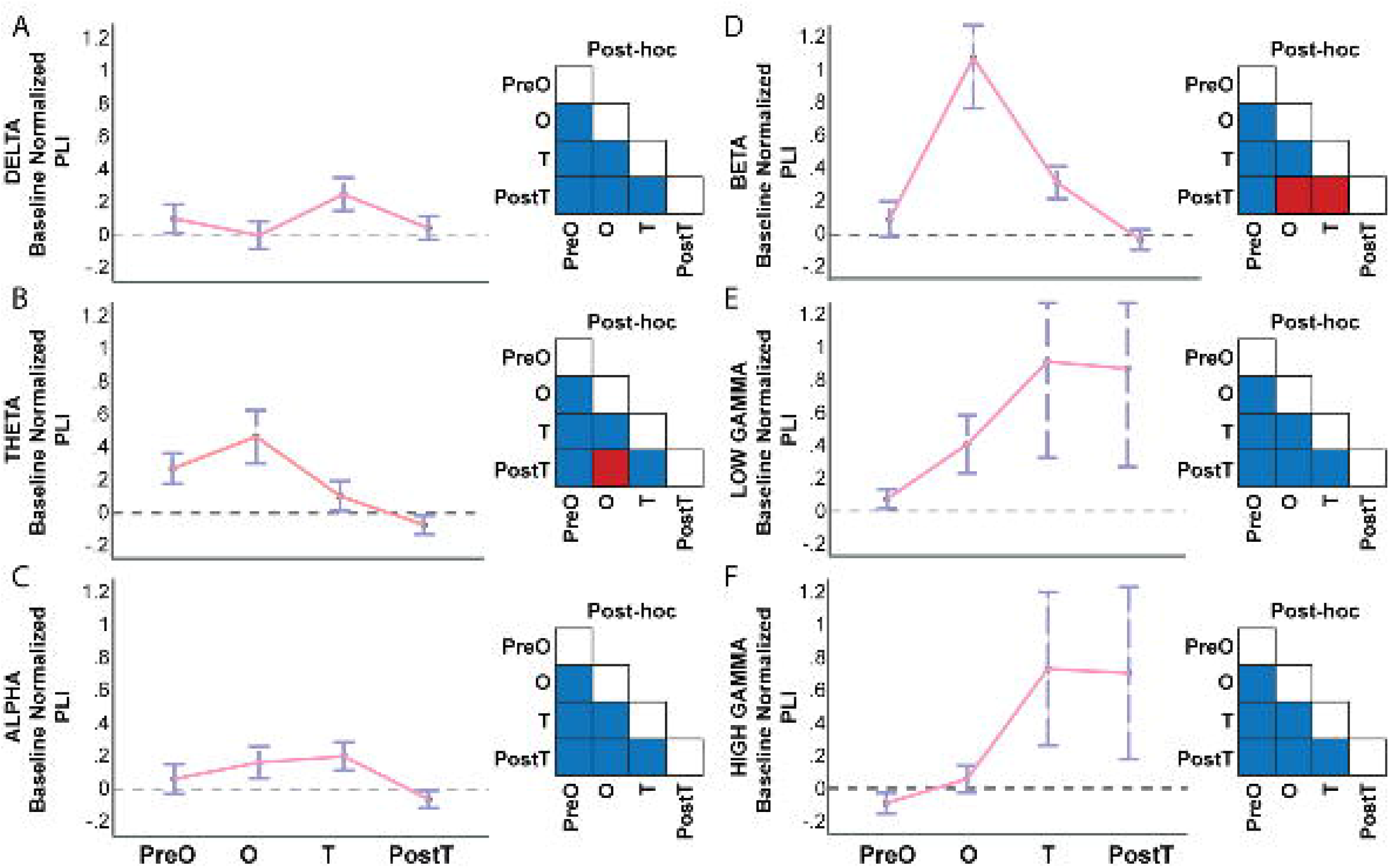
Thalamus to seizure onset zone phase synchronization (ANT-SOZ PLI) during seizure evolution: ANT-SOZ PLI focussed on the dynamic change of phase synchrony between ANT and SOZ alone. The PLI values were baseline normalized. Similar to global PLI, it was measured across the stages preonset (PreO), onset (O), Termination (T) and post-termination (PostT). (A-E) The line graphs with the error bars (mean±standard error) represent the change in phase synchrony across the different stages (preO to PostT) and frequencies (delta to high gamma). The corresponding lower-triangle insets represent inter-stage post hoc comparisons (Red – significant difference between stages and the blue no-significant difference between stages following Bonferroni correction).

### 3.5 Influence of disease burden on the thalamocortical synchronization

#### 3.5.1 ANT-ENA synchronization

Longer duration of epilepsy and increased seizure frequencies were associated with higher synchronization in the delta (stdβ=0.53, t=3.1, p=0.0035), theta, beta, and low gamma bands (stdβ’ s>0.395, t’ s>2.1, p’ s<0.046) at the seizure onset. At the termination, increased seizure frequencies and more prolonged duration were associated with higher delta (_std_β=0.58, t=3.5, p=0.002) and lower theta synchronization (_std_β=-0.97, t=-3.64, p=0.01).

#### 3.5.2 ANT-SOZ synchronization

Similarly, longer duration of epilepsy and increased seizure frequencies were associated with higher synchronization in the delta (_std_β=0.58, t=3.51, p=0.002) theta, beta, and low gamma bands (_std_β’ s>0.48, t’ s>2.7, p’ s<0.013) at the seizure onset. At seizure termination, increased seizure frequencies and more prolonged duration were associated with higher delta (≡tóβ=0.57, t=3.4, p=0.003) and lower theta synchronization (_std_β=-0.47, t=-2.59, p=0.016).

These results confirm that the disease burden affects ictal synchronization that extends beyond the seizure focus to subcortical structures.

### 3.6 ANT-SOZ phase-amplitude coupling at seizure onset and termination

At seizure onset, there was a significant increase in the coupling between the phases of low-frequency oscillations (delta, theta, and alpha) in the ANT and the amplitude of high gamma-band in the SOZ (Fig 6). At termination, there was a significant decrease in the coupling between the phases of low-frequency oscillations (delta, theta, and alpha) in the ANT and the amplitude of high gamma-band in the SOZ.

**Figure 6:**
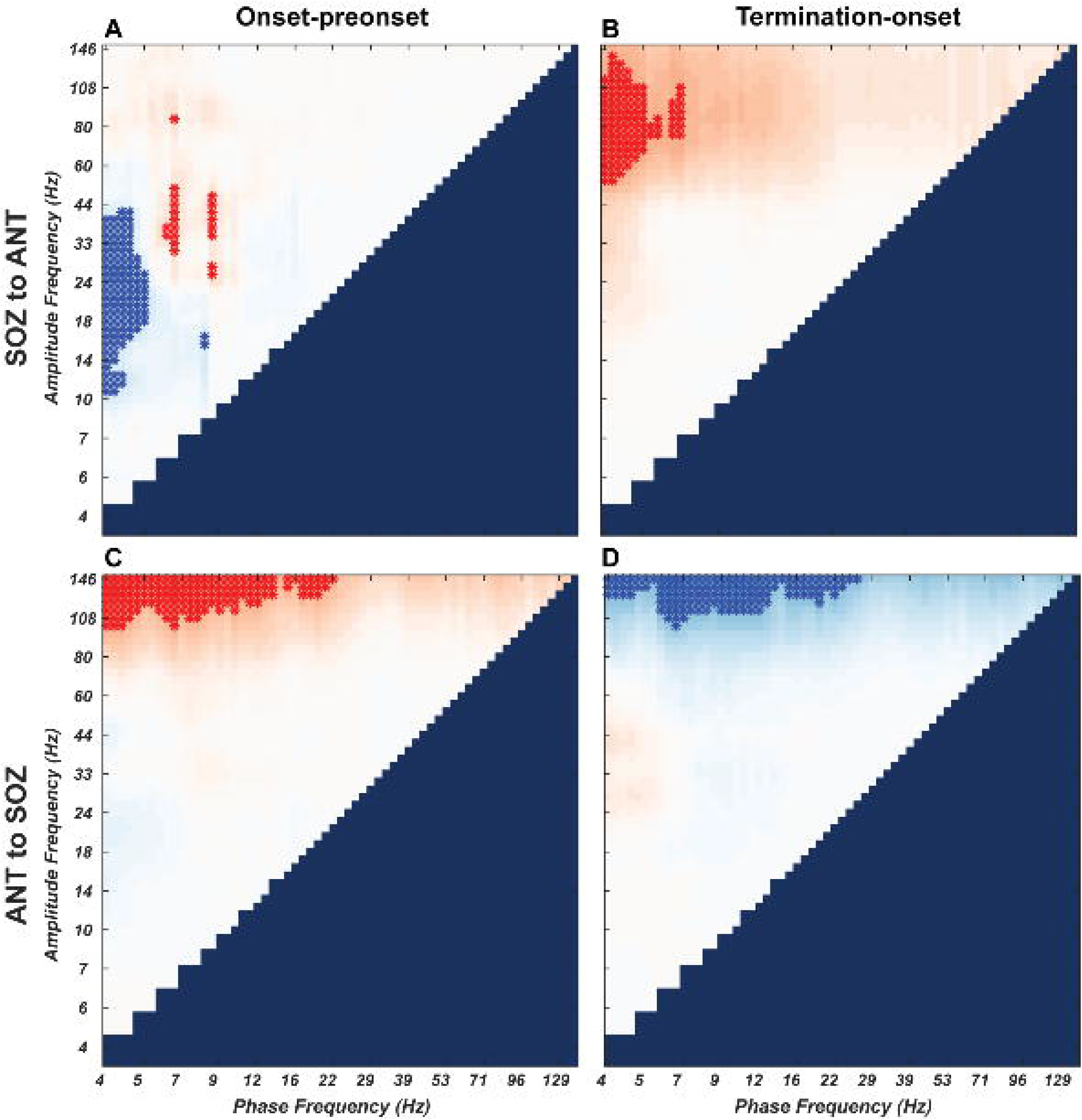
ANT-SOZ phase-amplitude coupling at seizure onset and termination: (A & C) These depict a bidirectional change in the phase-amplitude coupling between the seizure onset zone (SOZ) and the anterior nucleus of thalamus (ANT). The bidirectionality helps evaluate the coupling between the phase of low-frequency activity at SOZ to the amplitude of the high-frequency activity of ANT (A) and vice versa (C). We estimated the extent to which the modulation index (MI) matrix varies between pre-onset to seizure onset (A&C) while seizure onset to termination (B & D). Red asterisks indicate an increased coupling, and blue asterisks indicate a decreased coupling.

In contrast, when we tested how the phase of SOZ coupled with the amplitude of the ANT, we noted that at the onset, there was decreased coupling between the delta phase in SOZ with alpha-beta-low gamma amplitude in the ANT. Conversely, at the termination, there was a significant increase in the coupling between the delta phase in the SOZ and the amplitude of gamma-band in the ANT.

## 4. Discussion

Despite growing interest and a critical need in developing effective thalamocortical neuromodulation therapies, the mechanistic role of the thalamus in regulating seizures in TLE remains to be elucidated. We address this knowledge gap by capitalizing on the unique opportunity of LFP recordings directly from the human thalamus and its interconnected network during SEEG evaluation. The pertinent findings are: (1.) The ANT is coactivated at the onset of mesial temporal lobe seizures. (2.) Alteration in thalamocortical phase synchronization preceded seizure onset, and (3.) was higher at seizure onset. Finally, (4.) increased thalamic excitability paralleled increased thalamocortical coupling at the seizure onset, while termination was characterized by decreased thalamic excitability and decoupling. Based on these findings, we propose a coherent mechanism (Fig.7) that integrates local and long-range thalamic subnuclei specific neural interactions during seizures in drug-resistant nonlesional TLE.

**Figure 7:**
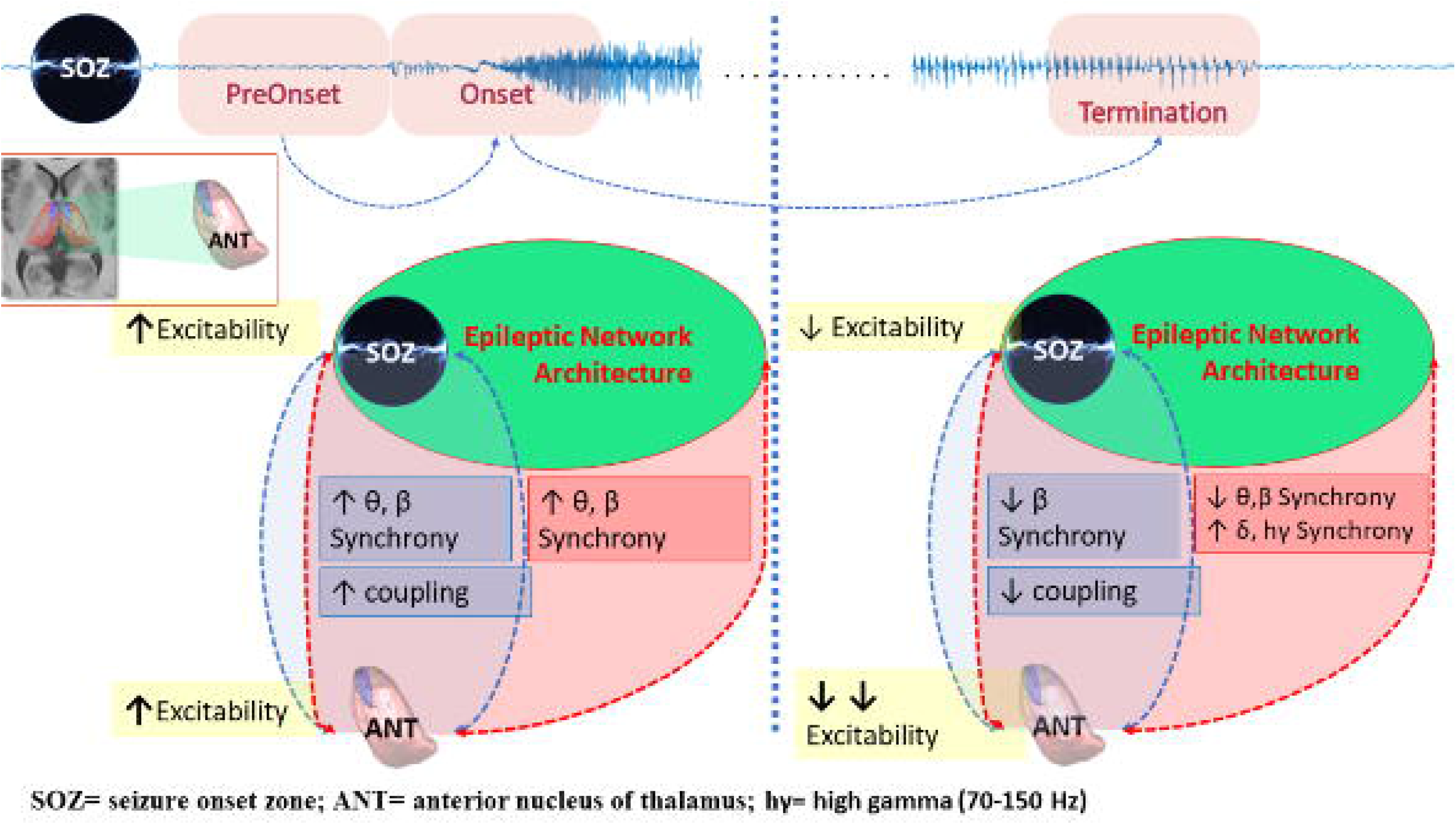
Graphical summary of thalamocortical network interactions during seizure onset and termination in temporal lobe epilepsy. Onset was characterized by increased network excitability, phaseamplitude coupling, and synchronization between the anterior nucleus of thalamus and cortical regions (including seizure onset zone and epileptic network architecture). The termination was characterized by decreased network excitability, decreased coupling, and synchronization.

### 4.1 Coactivation of ANT at the seizure onset

The increase in thalamic power spectra, phase synchronization, and increased PAC confirms the coactivation of the ANT at the onset of mesial temporal lobe seizures. The results are in agreement with preclinical studies demonstrating increased neuronal firing in the ANT at the onset of limbic seizures(Feng et al., 2017)(Sherdil et al., 2019). Limited electrophysiological studies in patients with focal epilepsy have also confirmed early ictal recruitment of the ANT(D. et al., 2018)(Osorio et al., 2015). In addition, ictal SPECT has consistently demonstrated increased blood flow in the thalamus prior to the generalization of focal onset seizures(Blumenfeld et al., 2009). Interestingly, our study is the first to demonstrate that even focal seizures that do not generalize can coactivate the ANT at the onset. Phases of slower oscillations are reported to be associated with increased neural excitability(Vanhatalo et al., 2004)(Alvarado-Rojas et al., 2011)(Breton et al., 2019). Similarly, our observation from power spectra, spectral exponents, and PAC suggests increased thalamic intrinsic excitability at seizure onset. The ANT being part of the Papez circuit receives afferents from the hippocampus via fornix and mammillary bodies through the mammillothalamic tract(Child and Benarroch, 2013). Seizures can propagate rapidly to ANT from the SOZ localized within the limbic network or indirectly after spreading to areas whose afferents back projects to the ANT.

### 4.2 Thalamocortical synchronization gates the progression of focal seizures

The thalamus can regulate intracortical synchronization and is hypothesized to be a “nodal hub” in the organization of epileptogenic networks in drug-refractory TLE(He et al., 2017). We observed a significant increase in thalamocortical synchronization from the baseline (i.g. interictal state) to the preonset state. Changes in synchronization and power spectra in the pre-seizure state have been well studied in cortical regions but remain to be explored in human thalamic subnuclei(Perucca et al., 2013)(Le Van Quyen et al., 2005). Such findings, if confirmed in a larger study, carry a translation significance, particularly in developing therapeutic feedback stimulation based on early detection or prediction. The thalamus is not a unitary structure, and likely different subnuclei will have a specific role in the seizure genesis and termination. The pulvinar subnuclei are proposed to facilitate seizure termination through increased synchronization while the mediodorsal and the ANT may facilitate the early organization of the seizure(Aracri et al., 2018)(Guye et al., 2006)(Evangelista et al., 2015)(Gale, 1992)(Cassidy and Gale, 1998). Indeed increased coherence at 5-20 Hz (theta and beta band) between the ANT and hippocampus was observed at the onset of induced seizures in experimental models of limbic epilepsy(Sherdil et al., 2019)(M.A. et al., 2003). Our study confirms the preclinical findings by using a non-linear measure of synchronization that is independent of the amplitude and less prone to volume conduction. Overall, the results confirm the dynamic engagement of ANT during ictogenesis in TLE. So what may be the role of thalamocortical synchronization during seizures? Below we propose a dynamic role of thalamocortical interactions during different stages of the seizure(Fig7).

LFPs recorded from the ANT represent a summation of synaptic inputs and spike firing(Buzsáki et al., 2012)(Manning et al., 2009). At seizure onset, the increased synchronization in lower frequencies (theta and beta) may be indicative of common incoming synaptic drive as higher frequencies tend to represent increased cell firing(Chrobok et al., 2018). It is well established that the number of corticothalamic inputs is greater than the number of thalamocortical fibers. Thus with increased recruitment during seizure progression, there is increased cortical feedback that modulates the thalamic intrinsic oscillatory state and firing patterns(Debay et al., 2001)(Bal et al., 2000). The two distinct firing modes of the thalamic cells are the tonic and bursting pattern(Murray Sherman, 2001). In chronic neurological illness, including TLE, pathologically higher bursting patterns were recorded from the ANT(Hodaie et al., 2006). The increased bursting patterns can cause thalamic hyperpolarization and the generation of low-frequency oscillations(Debay et al., 2001). Indeed, we observed a relative increase in the power of lower frequency oscillations (theta-delta) and an increased steepening of PSD towards seizure termination. Increased steepening of a spectral exponent decay has been associated with increased inhibition and dampening of propagation(Gao et al., 2017). Thus via activity-dependent changes in the thalamic intrinsic excitability and oscillations, the transmission of information (i.e., gating) within the reciprocal thalamocortical loop was regulated during a seizure. This was reflected in increased thalamocortical coupling at seizure onset, while decreased coupling towards seizure termination.

### 4.3 Clinical significance of studying the role of the thalamus in nonlesional epilepsies

Increasingly, epilepsy centers, including our level-IV center, are witnessing a higher number of nonlesional TLE. Unlike in “lesional” TLE (like TLE with hippocampal sclerosis-TLE-HS), where resection or ablation of the sclerotic hippocampus can significantly decrease or eliminate seizures, the surgical outcome in nonlesional TLE is considerably lower with pooled seizure freedom rates between 18-63%(Muhlhofer et al., 2017). The proposed hypothesis for the suboptimal surgical outcome in nonlesional TLE stems from emerging evidence that posits a difference in seizure circuitry between the two cohorts (nonlesional TLE vs. TLE-HS)(Reyes et al., 2016)(Vaughan et al., 2016). In TLE-HS, loss of hippocampal neurons from local excitotoxicity contributes to thalamic deafferentation and thalamic atrophy(Barron et al., 2014). The findings are often observed in clinical PET scans demonstrating thalamic hypometabolism(Jaisani et al., 2020). In contrast, in nonlesional TLE, it is hypothesized that recurrent chronic seizure spread to extratemporal regions, including the thalamus, causing remodeling and atrophy(Bonilha et al., 2010)(Riederer et al., 2008). In this context, our study confirms thalamic engagement even for focal seizures and the increased thalamocortical synchronization with a higher seizure burden and duration. The findings underscore the importance of treating seizures aggressively early in the disease course. The follow-up evaluation of treatment outcomes in our cohort was too short (less than a year) to make a meaningful association.

### 4.4 Study limitations

The study has multiple limitations. Due to ethical constraints with regard to thalamic implantation, our sample size was small and, more importantly, lacked a sufficient number of FBTC seizure types. Comparing the dynamics between focal vs. secondarily generalized seizures will be critical in exploring the role of ANT in the generalization of focal seizures(Blumenfeld et al., 2009). Another shortcoming in our analysis of functional connectivity between the thalamus and hippocampus was the lack of information about the direction of information flow (i.e., effective connectivity). Interestingly, one recent study explored the push-pull mechanism in focal epilepsy by studying directional information flow and suggested such a mechanism might be present between the thalamus and SOZ(Jiang et al., 2019).

## 5. Conclusion

The ANT mediates cortical-subcortical interactions between the limbic system and is hypothesized to facilitate the early organization of temporal lobe seizures. Using LFPs recorded directly from the limbic system, we confirm that the onset of mesial temporal lobe seizure is associated with increased thalamocortical network excitability and phase-amplitude coupling. Pre seizure state is associated with increased thalamocortical phase synchronization, thereby suggesting that the early organization of temporal lobe seizures involves the integration of the thalamocortical network. Towards seizure termination, there was a significant decrease in thalamic excitability, thalamocortical synchronization, and decoupling, thereby suggesting a breakdown in functional connectivity. A higher disease burden was significantly correlated with increased thalamocortical synchronization. Collectively, the results elucidate mechanistic insights and provide the temporal architecture of thalamocortical interactions that can be targeted in the rational designing of closed-loop interventions.

## ACKNOWLEDGMENTS

We would like to thank Miss. Auriana Irannejad for helping in data organization. GC, ET, DP, and SP would like to acknowledge the continuous support from the USA National Science Foundation Grant (NSF RII-2 FEC OIA-1632891) and NIH (1RF1MH117155-01).

## Abbreviations

ASD: Antiseizure drugs
EI: Epileptogenecity Index
ES: Electrographic Seizures
FAS: Focal Aware Seizures
FBTCS: Focal to bilateral tonic-clonic seizures
FIAS: Focal seizures with impaired awareness
LL: Line Length
PAC: Phase-amplitude coupling
PLI: Phase lag index
SOZ: seizure onset zone
SEEG: stereo-electroencephalography
TLE: Temporal lobe epilepsy
UEO: Unequivocal electrographic onset

